# Expected Genotype Quality and Diploidized Marker Data from Genotyping-by-Sequencing of *Urochloa* spp. Tetraploids

**DOI:** 10.1101/525618

**Authors:** Filipe Inácio Matias, Karem Guimarães Xavier Meireles, Sheila Tiemi Nagamatsu, Sanzio Carvalho Lima Barrios, Cacilda Borges do Valle, Marcelo Falsarella Carazzolle, Roberto Fritsche-Neto, Jeffrey B Endelman

## Abstract

Although genotyping-by-sequencing (GBS) is a well-established marker technology in diploids, the development of best practices for tetraploid species is a topic of current research. We determined the theoretical relationship between read depth and expected genotype quality (EGQ) for tetraploid vs. diploidized genotype calls. If the GBS method has 1% error, then 17 reads are needed to classify tetraploid samples as heterozygous vs. homozygous with 95% accuracy, compared with 63 reads to determine allele dosage. We developed an R script to convert tetraploid GBS data in Variant Call Format (VCF) into diploidized genotype calls and applied it to 267 interspecific hybrids of the tetraploid forage grass *Urochloa* (syn. *Brachiaria).* When reads were aligned to a mock reference genome created from GBS data of the *U. brizantha* cultivar ‘Marandu’, 25,678 bi-allelic SNPs were discovered, compared to approximately 3000 SNPs when aligning to the closest true reference genomes, *Setaria viridis* and *S. italica.* Crossvalidation revealed that missing genotypes were imputed by the Random Forest method with a median accuracy of 0.85, regardless of heterozygote frequency. Using the *Urochloa* spp. hybrids, we illustrated how filtering samples based only on GQ creates genotype bias; a depth threshold with corresponding EGQ equal to the GQ threshold is also needed, regardless of whether genotypes are called using a diploidized or allele dosage model.

## INTRODUCTION

*Urochloa* is the most cultivated genus as pasture on tropical livestock farms due to its tolerance to acidic soils, good carrying capacity, insect resistance, and nutritional value (Jank et al., 2014; Pessoa-Filho et al., 2017). The most economically important species are *U. decumbens* (syn. *Brachiaria decumbens)* and *U. brizantha* (syn. *B. brizantha*), which are both tetraploid (2n = 4x = 36). Apomixis is the normal mode of reproduction in these species, and for many years genetic improvement in South America was based on screening new introductions from Africa (Miles, 2007; Jank et al., 2011). To facilitate breeding by sexual hybridization, Swenne et al. (1981) utilized colchicine-induced tetraploids of the diploid species *U. ruziziensis* (2n = 2x = 18) as female parents to cross with apomictic tetraploids. This interspecific hybridization scheme has become the foundation of the *Urochloa* spp. breeding programs at CIAT and EMBRAPA (Lutts et al., 1991; Miles et al., 2006; Monteiro et al., 2016).

As in other crops, genome-wide markers can provide significant value for *Urochloa* spp. breeding programs. Several previous studies have utilized microsatellite markers to study population structure in *Urochloa* (Jungmann et al., 2010; Vigna et al., 2011; Silva et al., 2013), but the ubiquity and cost-effectiveness of SNPs are advantageous for discovering genetic variants and predicting complex traits. Arrays and genotyping-by-sequencing (GBS) of multiplexed, reduced-representation libraries have been utilized to generate large, bi-allelic SNP datasets in heterozygous tetraploids, including potato (Felcher et al., 2012; Uitdewilligen et al., 2013), alfalfa (Li et al., 2014), rose (Koning-Boucoiran et al., 2015), kiwi (Melo et al., 2016), and *Urochloa* spp. (Worthington et al. 2016; Ferreira et al. 2018). Both arrays and GBS generate a signal for each allele that can be used to predict allele dosage, i.e., the tetraploid genotype. For the SNP array, signal intensity is not necessarily proportional to allele dosage, and therefore different classification algorithms have been explored (Voorrips et al., 2011; Serang et al., 2012; Schmitz Carley et al., 2017).

For GBS data, the allele signal intensity is the read count, which can be analyzed using the aforementioned classifiers, but the focus of this manuscript is genotype calling based on a binomial model. The binomial model is central to the well-established software packages GATK (McKenna et al., 2010; Depristo et al., 2011) and FreeBayes (Garrison and Marth, 2012), as well as more recent tools developed specifically for polyploids (Blischak et al. 2018; Clark et al. 2018; Gerard et al. 2018). It is generally recognized that higher read depth is needed to estimate allele dosage in polyploids, but precise guidelines are lacking. Uitdewilligen et al. (2013) developed KASP assays for 270 GBS markers in potato and compared the genotype calls from the two methods; the results under different filtering criteria led the authors to conclude that “~60-80X can be used as a lower boundary for reliable assessment of allele copy number….” Bastien et al. (2018) used a threshold of 53 reads for determining allele dosage in potato because it was deemed “sufficient to distinguish between the five expected genotypic classes based on a chi-square distribution.”

Our first objective was to use probability theory to clarify the relationship between read depth and genotype quality (GQ) in tetraploids. GQ is a standard metric in the FORMAT field of the VCF file and defined as −10 log_10_(*q*), where *q* is the probability that the genotype call is erroneous (Danecek et al., 2011). Theoretical results were used to guide the analysis of GBS data for a panel of 267 *U. ruziziensis* x *U. brizantha* hybrids. Because few markers had sufficient read depth to determine allele dosage with reasonable accuracy, genotype calls were made using a diploid approximation, in which the three heterozygotes were not distinguished. This approximation is common for GBS in heterozygous tetraploids, and typically a threshold of 11 reads is used to ensure the probability of misclassifying a heterozygote as homozygous is less than 5% (Li et al., 2014). However, this threshold is based on the assumption of no error in the GBS method, and our theoretical treatment elucidates how the threshold increases with error.

Even with a diploid approximation, the *Urochloa* dataset contained missing data. Imputation of missing genotypes in GBS datasets has been studied extensively in inbred lines and heterozygous diploids, with Hidden Markov Models being the preferred method when a genetic or physical map for the markers is available (Hickey et al., 2012; Swarts et al., 2014; Fragoso et al., 2015). When a map is not available, as was the case for the *Urochloa* spp. hybrids, the Random Forest algorithm (Breiman, 2001) can still be used and has performed well in other species (Rutkoski et al., 2013; Money et al., 2015). Our objectives were to evaluate different filtering criteria, references genomes, and imputation accuracy in the tetraploid *Urochloa* dataset.

## MATERIALS AND METHODS

### Expected Genotype Quality

A binomial model was used to determine the statistical relationship between read depth and expected genotype quality (EGQ) for a particular genotypic class, such as ‘simplex’ for tetraploid genotypes or ‘heterozygous’ for diploidized genotypes. Let *f*(*k,N,ρ*) denote the probability mass function for the binomial distribution with *k* successes out of *N* trials and success probability *ρ*. The likelihood of observing *k* reads of the alternate allele given *N* total reads for tetraploid genotype *x* ∈ {0,1,2,3,4} was modeled as 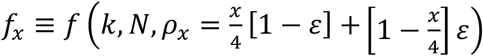, where the error rate *ε* is the probability that a read is generated by one allele but counted toward the other (e.g., due to errors during PCR or sequencing). Under a uniform prior, the maximum a posteriori (MAP) tetraploid genotype call for the observed result (*k,N*) is the value of *x* that maximizes *f_x_*. For some values of *k*, the MAP solution does not equal the true value. Summing *f* over these values of *k*, and expressing the result on the phred scale, leads to the following expression for EGQ_tet_:

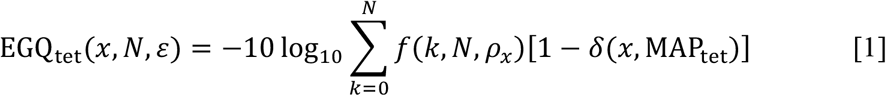

The symbol *δ* in Eq. 1 is the Kronecker delta function, which equals 1 when its two arguments are equal and 0 when they are unequal. (While this manuscript was in preparation, Gerard et al. (2018) independently published a result similar to Eq. 1 called the “oracle misclassification error rate.”)

For diploidized genotype calls, the three possible genotypic states are denoted {*A,H,B*}, where the heterozygous state *H* = dosages 1, 2, or 3, and the homozygous states *A* = dosage 0 and *B* = dosage 4. The corresponding 3-vector of posterior probabilities is proportional to (*p_A_,p_H_,p_B_*) ≡ (*f*_0_,*f*_1_ + *f*_2_ + *f*_3_,*f*_4_), and the MAP solution for the observed result (*k,N*) is the value of *j* that maximizes *p*,·. For some values of *k*, the MAP solution does not equal the diploidized genotype *y* corresponding to the true tetraploid state *x*. Summing *f* over these values of *k*, and expressing the result on the phred scale, leads to the following expression for EGQ_dip_:

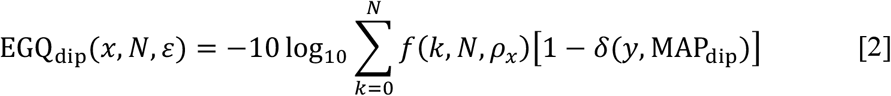

Although Eq. 1 and 2 tend to increase with read depth, they are not monotone functions of *N*. Our results for EGQ correspond to the monotone extension

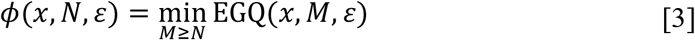

which has the property *ϕ*(*x, N, ε*) ≥ *ϕ*(*x, M, ε*) for *N* > *M*. Using the R programming language (R Development Core Team, 2017), a function was created (Supplemental File S1) to calculate *ϕ*(*x, N, ε*).

### GBS of *Urochloa* spp

Genomic DNA was extracted using the Qiagen DNeasy kit for 267 tetraploid *U. ruziziensis* x *U. brizantha* hybrids from Embrapa Beef Cattle, as well as for the *U. brizantha* cultivar ‘Marandu’. GBS libraries were prepared according to Elshire et al. (2011), using the ApeKI enzyme and sequenced on five lanes of the Illumina Hi-Seq 2500 platform with 1×100 bp reads. Reads were demultiplexed and trimmed using Cutadapt (Martin, 2011) and then aligned to five different *Poaceae* genomes with bwa-mem (Li, 2013): *Setaria viridis* (DOE-JGI, 2018a), *Setaria italica* (Bennetzen et al., 2012), *Sorghum bicolor* (DOE-JGI, 2018b), *Oryza sativa* (Ouyang et al., 2006), and *Zea mays* (Schnable et al., 2009). The alignment percentage for each reference was evaluated with Bowtie2 (Langmead and Salzberg, 2012). Reads were also aligned to a *U. brizantha* mock reference genome generated from the reads for ‘Marandu’ with the GBS-SNP-CROP pipeline (Melo et al., 2016). The Genome Analysis Toolkit (GATK, McKenna et al., 2010; Depristo et al., 2011) HaplotypeCaller was used for SNP discovery, followed by removal of SNPs that did not meet the recommended thresholds (GATK, 2016): FS (Fisher Strand Bias) ≤ 60.0, MQ (RMS Mapping Quality) ≥ 40.0, MQRankSum (Rank Sum Test for Mapping Quality) ≥ −12.5, ReadPosRankSum (Rank Sum Test for Read Position) ≥ −8.0.

Using the R programming language, a function was created (readVCF, Supplemental File S2) to process the VCF file and perform additional filtering. Only bi-allelic SNPs were retained. The VCF file includes variants relative to the reference genome, regardless of whether they are polymorphic in the genotyped population. To identify polymorphic markers, the total number of reads for the minor allele, or minor allele depth (MAD), was calculated for each marker based on the AD field, and variants with MAD < 2 were removed. GATK calculates allele frequency based on the dosage of called genotypes, which was deemed unreliable due to low read depth. A suitable proxy for filtering that does not require allele dosage information is the frequency of genotypes homozygous for the major allele (HMA), which was capped at 0.99. For each sample, GATK provides the phred-scaled likelihood (PL) for each of the 5 tetraploid genotypes, which was converted into a posterior probability *p_i_* for genotype *i* ∈ {0,1,2,3,4} (assuming a uniform prior) by

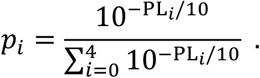

The tetraploid genotype call corresponds to the largest probability, and 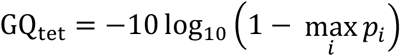.

Due to the low read depth per sample in the *Urochloa* dataset, diploidized genotype calls were made in which the three heterozygous genotypes were not differentiated. This corresponds to defining a new vector of posterior probabilities, 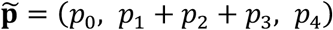, in which the probability of the heterozygous state is the sum of the probabilities for the simplex, duplex, and triplex genotypes. The diploidized genotype call corresponds to the largest probability, and 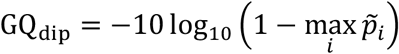.

Missing genotypes were imputed with the R package randomForest (Liaw and Wiener, 2002; Supplemental File S3), which is based on the algorithms in Breiman (2001). For each marker, a training set of 100 clones was randomly selected from the clones with genotypes, and all other clones with genotype data were used for validation. Because each marker had no more than 50% missing data, this ensured at least 33 clones were available for validation. 300 classification trees were used for prediction, and all markers with *r*^2^ ≥ 0.1 were used as *m* potential predictors. We used the default setting of randomly sampling 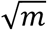 predictors at each split. Classification accuracy is the proportion of clones in the validation set for which the predicted genotype is correct.

## RESULTS

### Expected Genotype Quality

A binomial model was used to determine the statistical relationship between read depth and expected genotype quality (EGQ) for a particular genotypic class, such as ‘simplex’ for tetraploid genotypes or ‘heterozygous’ for diploidized genotypes. EGQ involves the expectation over all possible allele counts at a particular depth, whereas GQ corresponds to a particular allele count. In addition to read depth, the other key parameter affecting EGQ is the error rate, defined as the probability that a read is generated by one allele but counted toward the other (e.g., due to errors during PCR or sequencing). Since EGQ is reported on the phred scale, a score of 13 corresponds to 95% accuracy, and a score of 20 corresponds to 99% accuracy.

Figure 1 shows how EGQ differs for simplex vs. duplex genotypes, as well as when allelic dosage is estimated (blue) vs. diploidized calls (green). Higher accuracy is achieved for simplex compared to duplex samples when allelic dosage is determined, but under diploidized genotype calling the reverse is true. The intuitive reason for this result is that a duplex genotype can appear as either simplex or triplex due to sampling variation, but comparable uncertainty for the simplex genotype exists only in the direction of higher dosage (i.e., with the duplex). If dosage is not determined, however, then simplex genotypes are more readily confounded with nulliplex homozygotes than duplex samples are with either homozygote. In the absence of error (solid lines), 11 reads are needed to make diploidized genotype calls with 95% accuracy, compared with 61 reads for tetraploid genotypes. As Figure 1 shows, allelic errors have a greater effect on EGQdip than EGQtet. With 1% error (dashed lines), the minimum depth needed to achieve 95% accuracy for diploidized genotypes increases to 17 reads, while the minimum depth for tetraploid genotypes increases to 63 reads.

**Figure 1.**
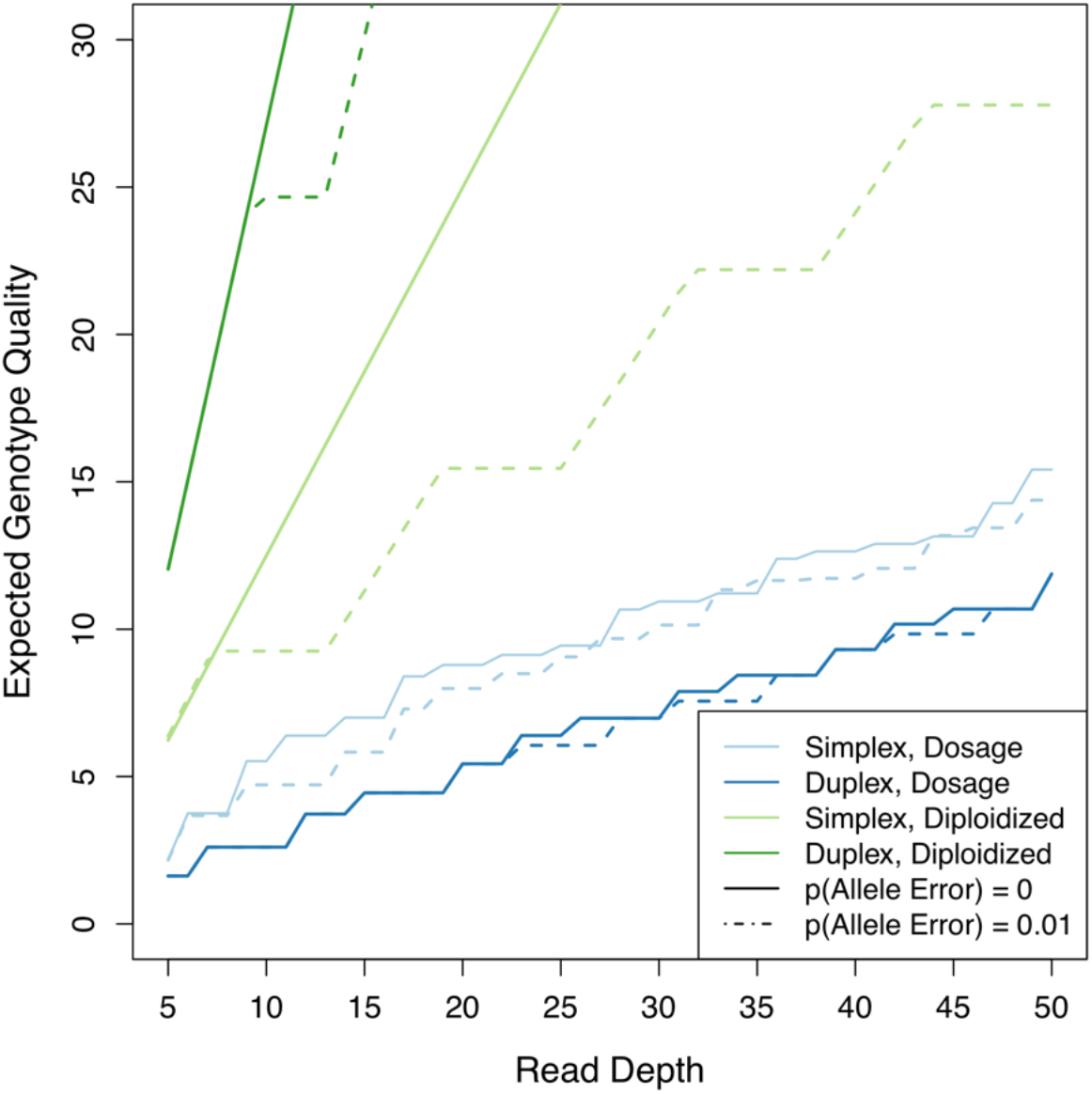
Expected Genotype Quality (EGQ) as a function of read depth, for two different allele error rates.

### GBS of *Urochloa* spp. hybrids

As no reference genome for the *Urochloa* spp. hybrids was available, the reference genomes of five other *Poaceae* species were evaluated for alignment. Figure 2 shows the number and percentage of aligned reads from the ApeKI-reduced representation of the *U. brizantha* cultivar ‘Marandu.’ The percentage of reads aligned was low for all genomes, ranging from 1.92% for *Oryza sativa* to 7.88% for *Setaria italica.* For both *Setaria* species and *Sorghum bicolor,* over 3/4 of the aligned reads mapped to a unique location. For *Oryza sativa* and *Zea mays,* this proportion decreased to 1/2. The same five genomes were compared with respect to variant discovery in a panel of 267 tetraploid *U. ruziziensis* x *U. brizantha* hybrids. After removing variants with median depth < 8, the two *Setaria* species generated the most bi-allelic SNPs, in the range 2809−3203 (Table 1).

**Figure 2.**
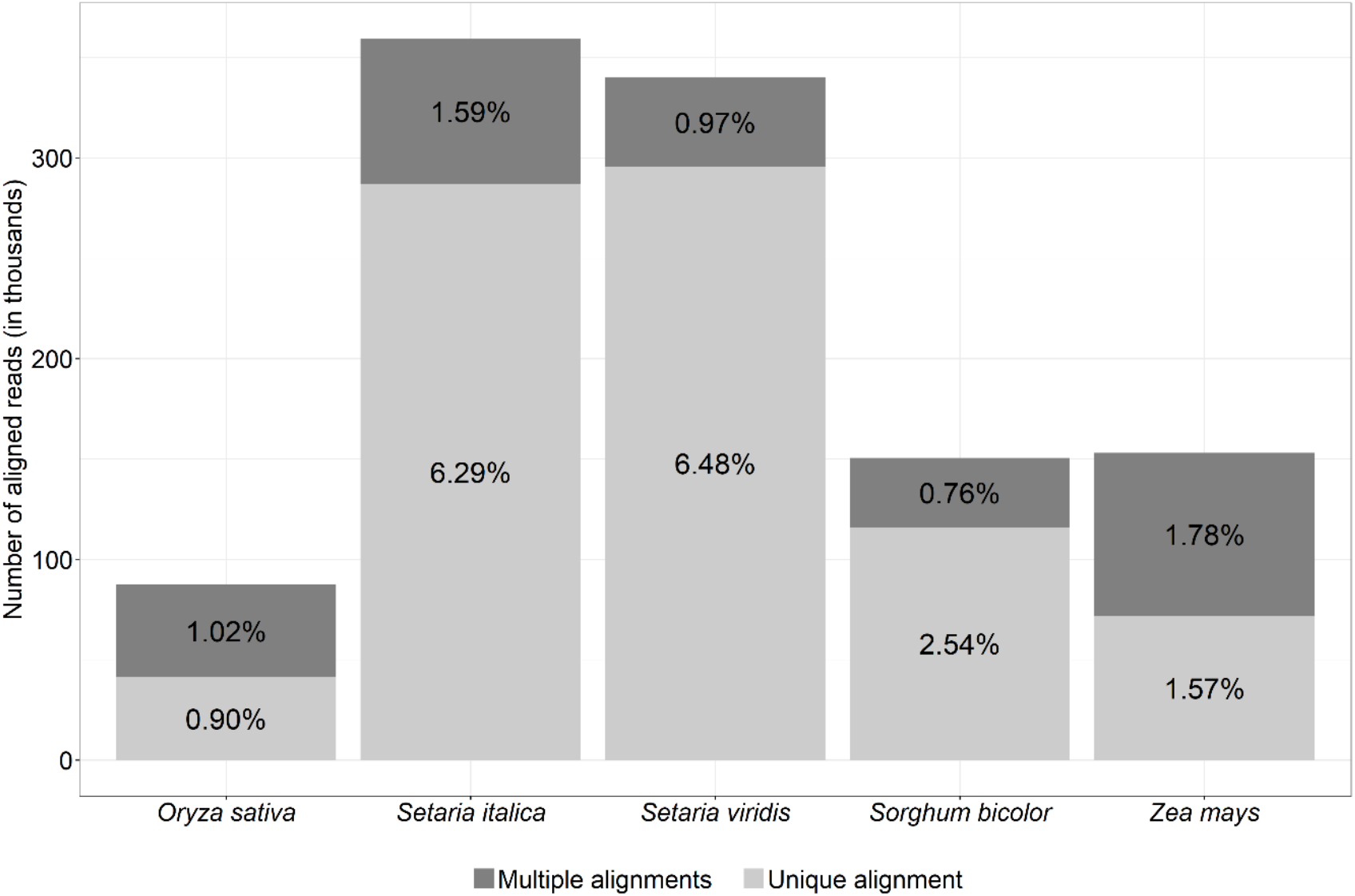
Number and percentage of reads from the *U. brizantha* cultivar ‘Marandu’ that aligned to five *Poaceae* reference genomes.

**Table 1.** Number of bi-allelic SNPs with < 50% missing data based on a minimum sample depth of 8.

| Reference | # SNPs |
| --- | --- |
| *U. brizantha* | 25,678 |
| *S. viridis* | 3,203 |
| *S. italica* | 2,809 |
| *S. bicolor* | 1,331 |
| *Z. mays* | 763 |
| *O. sativa* | 571 |

To better utilize the GBS reads, a mock reference genome was built by clustering the trimmed reads from ‘Marandu’ into 1,309,910 non-redundant, consensus sequences, or “centroids” (Melo et al., 2016). A highly repetitive sequence was detected in the centroids, for which the first 50 bp are GAGATCGGAAGAGCGGTTCAGCAGGAATGCCGAGACCGATCTCGTATGCC.

The entire 50 bp was present in 3.3% of the centroids, and when truncated to the first 40 or 30 bp, the frequency increased to 8.5% and 14.9%, respectively. The repetitive sequence was also detected in all 267 hybrids. A nucleotide BLAST search of the 50 bp sequence against the NCBI database returned highly significant matches to a diverse set of species, including *Larimichthys crocea* and *Cyprinus carpio* (100% identity across 49 bp), *Triticum aestivum* and *Solanum pennellii* (98% identity across 50 bp).

When the GBS reads for the 267 hybrids were aligned to the centroids, the number of bi-allelic SNPs with median depth ≥ 8 increased to 25,678 (Table 1). According to the binomial model, a depth threshold of 8 reads corresponds to EGQdip = 10 for simplex genotypes (assuming no GBS error). Figure 3 is a histogram of the GQ scores for all 153,589 genotypes (sample x marker combinations) with depth = 8 in the filtered dataset. The peak at GQdip = 10 for homozygous genotypes illustrates how EGQdip of the simplex (or triplex) genotype constrains GQ for homozygotes. By contrast, the heterozygous genotype calls with depth = 8 have GQ scores over 30. According to the binomial model, a depth threshold of 42 is needed for EGQ_tet_ ≥ 10 (assuming no GBS error). As only 2,338 SNPs had median depth ≥ 42, tetraploid genotype calls were not pursued.

**Figure 3.**
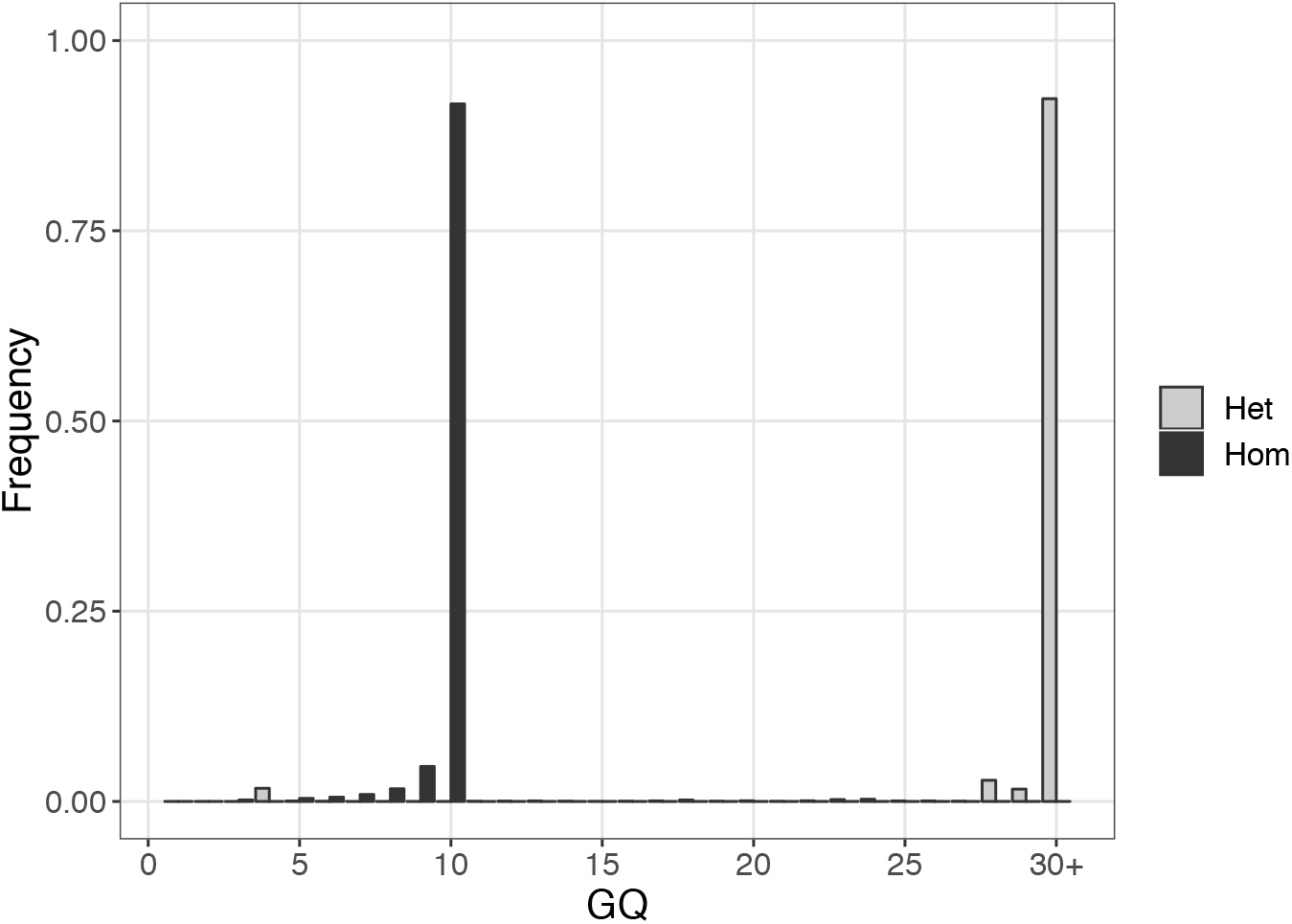
Distribution of genotype quality (GQdip) scores for diploidized genotypes with sample depth = 8 in a filtered set of 25,678 SNPs, discovered using the *U. brizantha* mock reference genome for alignment. Heterozygous samples (“Het”) are shown in light gray, and homozygous samples (“Hom”) are shown in dark gray.

The cumulative distribution in Figure 4 reveals the SNP dataset is dominated by rare alleles. The x-axis of Fig. 4 is the genotype frequency of clones homozygous for the major allele (HMA), and the y-axis is the proportion of SNPs for which the HMA frequency is less than or equal to the x-axis value. The SNP counts in Table 1 are based on an upper limit of 0.99 for HMA, and 51% of the SNPs discovered with the mock reference genome had HMA genotype frequencies between 0.95 and 0.99.

**Figure 4.**
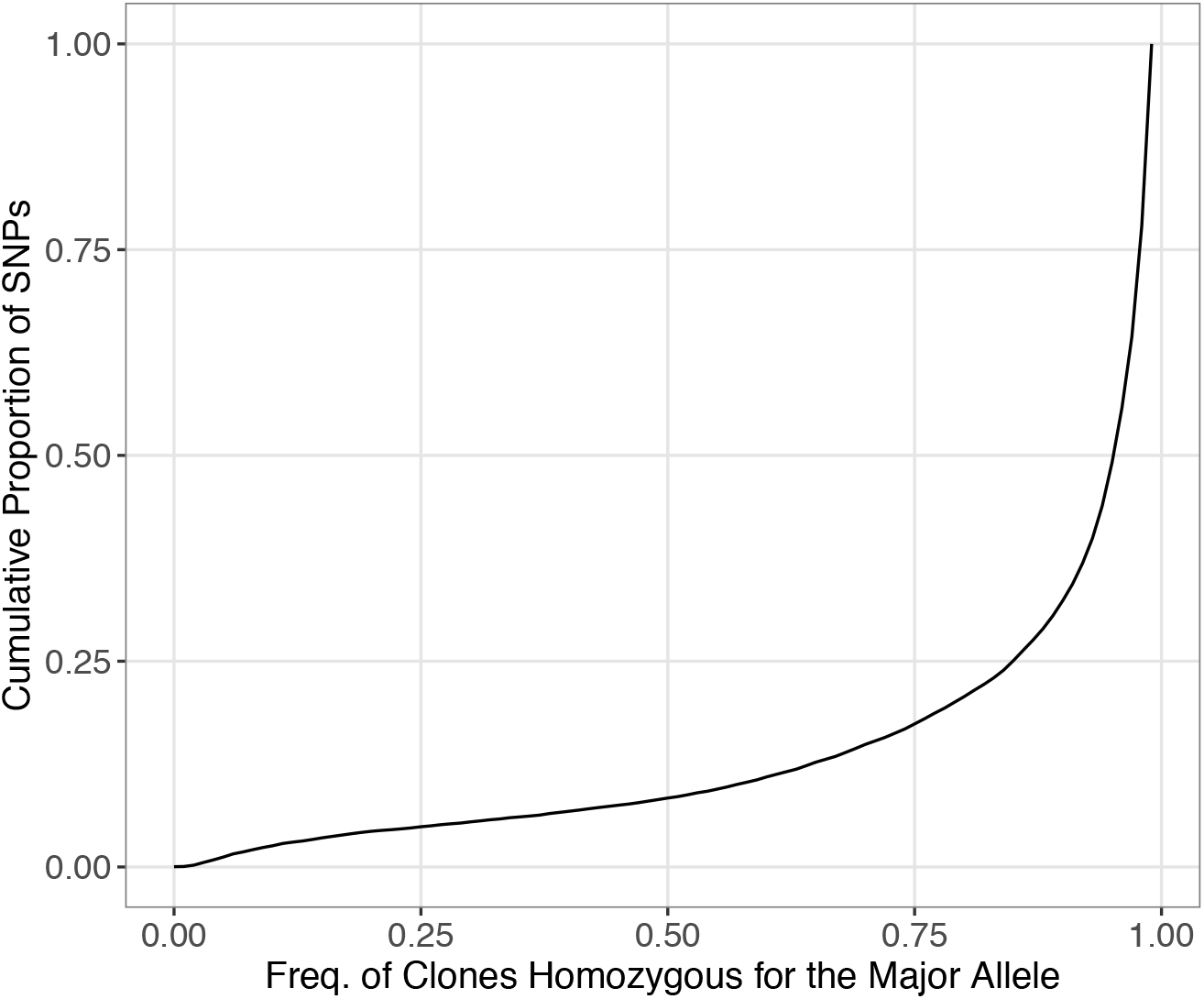
Cumulative distribution for the genotype frequency of clones homozygous for the major allele (HMA), based on 25,678 bi-allelic SNPs discovered using the *U. brizantha* mock reference genome for alignment. The y-axis is the proportion of SNPs for which the HMA frequency is less than or equal to the x-axis value.

### Genotype Imputation

The success of genotype imputation depends on the amount of linkage disequilibrium (LD) between markers, which is often quantified by the physical distance at which *r*^2^ (the squared correlation) drops below some threshold. Since a physical reference genome was unavailable for this study, LD was quantified based on the maximum *r*^2^ for each SNP. Figure 5A shows the distribution of *r*^2^max for the 3,230 SNPs from the filtered dataset that are 25–75% heterozygous, to capture a range of difficulty for imputation. The median value of *r*^2^_max_ was 0.4–0.5 for heterozygote frequencies below 0.5 but gradually decreased as the proportion of heterozygotes increased toward 0.75.

**Figure 5.**
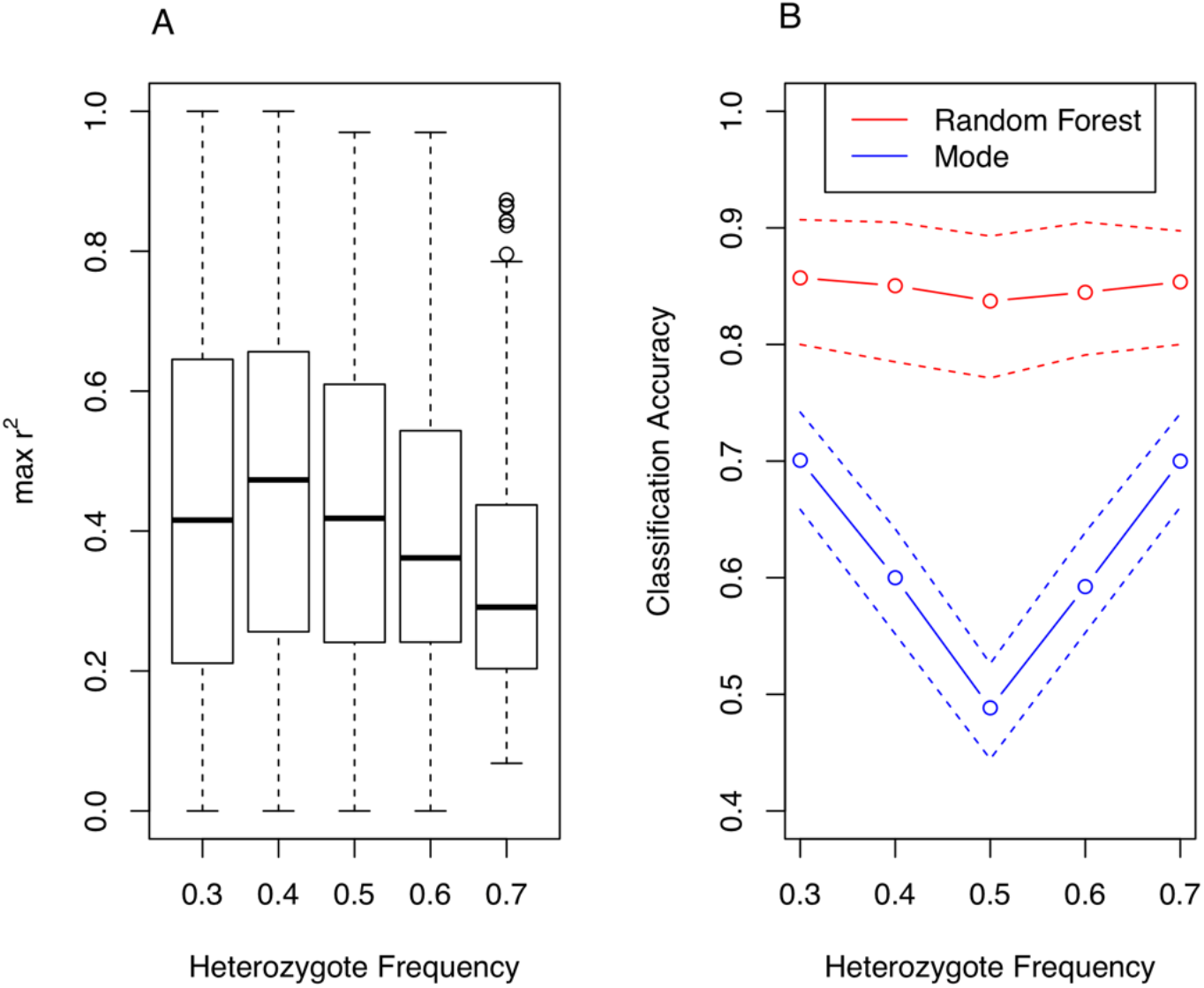
(A) Distribution of the maximum LD (*r*^2^) for each marker, binned by heterozygote frequency. (B) Imputation accuracy, defined as the proportion of imputed values equal to the masked value.

Cross-validation accuracy was determined with a training set of 100 clones, selected at random from all clones with genotype data for a particular marker. The accuracy shown in Figure 5B is the proportion of predicted values equal to the masked value. The results are binned by heterozygote frequency, with the median accuracy shown by a solid line and the first and third quartiles by dashed lines. Imputation with the population mode is a simple baseline method that, by definition, has lower accuracy as the frequency of the modal genotype declines. By contrast, the Random Forest method was largely unaffected by heterozygote frequency, with a median accuracy of approximately 0.85.

## DISCUSSION

As mentioned in the introduction, there has been variation in the filtering criteria used in previous studies involving GBS of tetraploids. One of the most cited is Uitdewilligen et al. (2013), who recommended 60–80X to determine allele dosage. Our theoretical calculations indicate this range corresponds to 95–98% genotype accuracy for GBS error rates below 1%. For diploidized genotype calling, the threshold of 11 reads in Li et al. (2014) has been commonly used by others, which corresponds to 95.8% genotype accuracy in the absence of error but only 88.1% genotype accuracy when the GBS error is 1%. To achieve 95% genotype accuracy with 1% GBS error, 17 reads are needed, and 98% genotype accuracy requires 27 reads.

The need for higher read depth per site to make accurate genotype calls in tetraploids underscores the importance of selecting restriction enzymes to optimize the fragment size distribution. This study utilized ApeKI, which has a 5 bp recognition sequence, while Worthington et al. (2016) and Ferreira et al. (2018) used enzymes with 6 bp recognition sequences (HincII and NsiI, respectively) for GBS of *Urochloa* spp. F1 populations. Future research on GBS for *Urochloa* should explore a two enzyme-system, such as the PstI/MspI combination introduced by Poland et al. (2012) for barley and wheat, as a way of generating more markers with higher read depth. Bastien et al. (2018) compared ApeKI against PstI/MspI in tetraploid potato and obtained tenfold more markers with the two-enzyme system when using a minimum sample depth of 53 reads.

The difference in expected genotype quality for simplex (or triplex) vs. duplex genotypes has important implications for filtering GBS data. Setting a minimum GQ value will create bias against duplex samples when calling tetraploid genotypes, and against simplex/triplex genotypes with diploidized genotypes. Using a depth threshold corresponding to the desired minimum EGQ does not introduce this bias, but this does not address the issue of reads with low base or mapping quality. A combination of the two approaches seems best, using a depth threshold with EGQ equal to the GQ threshold. Supplemental File S1 can be used to calculate EGQ (Eq. 3) for any depth and error rate, and Supplemental File S2 can be used to generate matrices (marker x sample) of tetraploid or diploidized genotype calls and corresponding GQ scores from a VCF file.

The aforementioned considerations are appropriate for genotype calling based on the posterior mode. An alternative approach is to estimate allele dosage based on the posterior mean, which produces fractional genotype calls (Ashraf et al., 2014; Sverrisdóttir et al., 2017). Such data are suitable when additive models are used in association analysis and genome-wide prediction, but a number of genetic analyses require integral estimates of dosage, including linkage analysis (Hackett et al., 2013; Zheng et al., 2016), dominance effects (Rosyara et al., 2016; Endelman et al., 2018), and haplotype inference (Su et al., 2008; Aguiar and Istrail, 2013).

This study used the traditional approach of setting hard thresholds for genotype calling, followed by imputation of the missing data. We used depth and GQ thresholds to achieve 90% genotype accuracy and allowed for up to 50% missing data per marker, which were imputed with accuracy close to this value (75% of the samples had 80–90% accuracy). We did not explore the interplay between GQ threshold and imputation accuracy, but this is an interesting topic for future research. It seems appealing to select thresholds to achieve similar accuracy in the samples called based on allele counts vs. those that are imputed. Ultimately, the traditional two-step approach (threshold then impute) is suboptimal because the read counts for the missing genotypes are not utilized during imputation. For ordered markers, this limitation can be overcome by using Hidden Markov Models (HMMs) with read counts as the emission states. This approach has been used in diploid mapping populations (Fragoso et al., 2015; Bilton et al., 2018) and can be extended to the HMMs developed for SNP array data in tetraploids (Hackett et al., 2013; Zheng et al., 2016). For unordered markers, alternative imputation methods need to be explored.

## Acknowledgments

Financial support was provided by the National Council for Scientific and Technological Development (CNPq), Coordination for the Improvement of Higher Education Personnel (CAPES), and the Brazilian Agricultural Research Corporation (EMBRAPA). Computing services were provided by the National Center for High Performance Processing in São Paulo (CENAPAD) and the Center for High Throughput Computing (CHTC) at UW-Madison. We thank Schuyler Smith for contributions to the bioinformatics pipeline and Sushan Ru for comments on the binomial model.

## Author contributions

JBE and FIM designed the study. SCLB and CBdoV crossed and developed the *Urochloa* population. KGXM performed the DNA extraction. STN and MFC built the mock reference genome. JBE, FIM and STN analyzed the data and drafted the manuscript. RFN and MFC provided analytical expertise and edited the manuscript. JBE and RFN supervised the whole study. All authors read and approved the final version of the manuscript for publication.

